# A Neuroendocrine Circuit That Suppresses Excretion and Pelvic Pain During Activity

**DOI:** 10.64898/2026.07.23.736864

**Authors:** Lucas E. Cabrera-Zapata, Archana Venkataraman, Andrea M. Harrington, Jitao Hu, Fernanda Castro-Navarro, Kristina Galatsis, Xin Duan, Stuart M. Brierley, Kimberly Keil Stietz, Holly A. Ingraham

## Abstract

Mate-seeking during peak sexual receptivity is tightly coupled with exploratory movement, a process made more efficient by transient suppression of excretion. Here, we define a functional hypothalamic-hindbrain circuit in mice that promotes locomotion while eliminating urination and defecation for hours. Chemogenetic stimulation of excitatory estrogen-melanocortin-responsive MC4R^+^ neurons in the ventrolateral ventromedial hypothalamus (VMHvl) effectively silences bladder and colonic visceral reflexes, even at noxious distension pressures, underscoring the potency of this anti-excretion circuit. Pelvic sensations and excretion are restored only after ablating inhibitory GABAergic neurons in the Barrington’s nucleus/locus coeruleus (BAR/LC) hindbrain region or after antagonizing endogenous endorphin signaling. Our study illustrates how a hormone-responsive brain node prioritizes movement over excretion and blunts pelvic discomfort, thereby optimizing an essential voluntary behavior for evolutionary fitness.

## MAIN TEXT

The brain’s ability to prioritize voluntary behaviors is essential for species propagation and self-survival. Of these, initiating and sustaining exploratory behavior for seeking mates or resources requires goal-directed locomotion. However, this intentional choice to move suppresses other voluntary behaviors, including urination and defecation, as controlling when and where to excrete is critical when time is limited or when avoiding predators.

Neural control of excretion has been studied primarily in the context of urination and involves multiple brain regions and an intricate set of ascending and descending spinal pathways within the peripheral nervous system, as reviewed in (1, 2). The primary brain region of focus for urination is Barrington’s nucleus (BAR), located medial to the locus coeruleus (LC) below the fourth ventricle. For more than a century, it has been known that voluntary control of bladder function depends on the BAR, also known as the pontine micturition center (PMC) (3). Among the diverse subsets of neurons within BAR, as recently defined by single-cell and spatial transcriptomics (4, 5), glutamatergic excitatory neurons are essential for voiding, as optogenetic stimulation of BAR^VGLUT2^ neurons triggers micturition, whereas their ablation causes urinary retention (6). Many of these excitatory neurons in BAR express the neuropeptide corticotropin-releasing hormone (CRH). Multiple groups have shown that BAR^CRH^ neurons descend to the sacral spinal cord parasympathetic nuclei (SPN) containing preganglionic bladder motor neurons and promote micturition, as well as male patterning of urine marks (7). Similarly, two other subsets of excitatory neurons within BAR expressing *Penk* (BAR^PENK^) or *Esr1* (BAR^ESR1^), relax the urethral sphincter and promote voluntary or context-dependent scent marking in male mice (5, 8). While far less is known about the role of BAR in defecation, isolated reports have linked both BAR and LC to colonic activity, as both of these brain regions receive ascending spinal inputs from the colon (9-11). Using chemogenetics, Ogawa and colleagues recently established a functional relationship between BAR/LC^VGLUT2^ neurons and fecal output (12), expanding the role of the BAR/LC as a defecation brain center. Although these collective studies address how neural circuits promote excretion, it remains unclear whether dedicated circuits exist that inhibit hind-brain-spinal cord reflexes and temporarily block voiding, especially when other volitional behaviors are prioritized.

Within the hypothalamus, neurons in the VMHvl regulate several social-behavioral outputs, including male aggression, conditioned fear (13-16), female social behaviors (17), and maternal aggression (18). VMHvl neurons are marked by *Esr1*, encoding estrogen receptor alpha (ERα), but lack expression of the other prominent VMH marker, *Sf-1* (19). A subset of these VMHvl^ESR1^ neurons is highly sensitive to estrogen and is distinguished by hormone-dependent upregulation of the melanocortin 4 receptor (MC4R) (20). Convergence of these two pathways in VMHvl^ESR1/MC4R^ neurons increases voluntary movement (21), driving the pre-ovulatory activity spike in female mammals, as originally observed over a century ago (22).

Our earlier studies, along with others using monosynaptic anterograde tracers, showed that, among the many projections originating from estrogen-sensitive VMHvl^MC4R^ neurons, strong connections are observed to the arousal center in the dorsal medial pontine region, near the LC, with faint projections to BAR (20, 23). However, in contrast to other brain regions that project to BAR, including the lateral hypothalamic area (LHA), medial preoptic area (MPOA), and the midbrain periaqueductal grey (PAG) (6, 7, 24-26), a functional connection between the VMHvl and bladder or colon has yet to be established. Here, using a newer-generation anterograde transsynaptic tracer (27), we identified strong VMHvl^MC4R^ afferents to BAR/LC and asked whether they coordinate a broader set of voluntary behaviors beyond physical activity, including excretion. These new findings reveal a powerful neuroendocrine-brainstem circuit that prioritizes physical activity while suppressing colonic and bladder spinal-brainstem reflexes, effectively stopping all excretion and silencing visceral discomfort.

### ESR1/MC4R Neurons in the VMHvl Project to the Hindbrain and Suppress Micturition

Prior work mapping anterograde projections of VMHvl^ESR1^ neurons noted caudal innervation of midbrain and brainstem regions, including the ventrolateral periaqueductal grey (vlPAG) and the LC (20). Using an AAV2 Cre-dependent anterograde transsynaptic mWGA-mCherry viral tracer (mWmC) (Figures 1A and 1B), we found strong mCherry^+^ signals marking post-synaptic neurons in BAR from the VMHvl^MC4R^. Other projections included those to tyrosine hydroxylase (TH) noradrenergic LC neurons adjacent to BAR and to the vlPAG in the midbrain (Figures 1C and S1). This projection pattern was observed in male (N = 4) and female (N = 5) brains, prompting us to ask if chemogenetic activation of VMHvl^MC4R^ neurons using Cre-dependent DREADDs (Designer Receptors Exclusively Activated by Designer Drugs; AAV2-DIO-hM3Dq-mCherry, sDREADDs) injected into the VMHvl of *Mc4r*^*Cre/+*^ mice might affect micturition. We first validated the void spot assay (VSA) in our hands (28, 29 and Figure 1D), and confirmed that the high-affinity, selective agonist deschloroclozapine (DCZ, 100 µg/kg) (30) does not alter the stereotypic corner preference or the quantity of urine released in sham or control mice (Figure 1E). Stimulating VMHvl^MC4R^ neurons eliminated all urination and fecal pellets for at least 2 hours while simultaneously increasing locomotor activity with no evidence of overt aggression (Figures 1F, Figure S2A, S2B, and Movies S1 and S2), whereas inhibiting VMHvl^MC4R^ neurons promoted urination and suppressed activity (Figures 1G and S2C). Another hypothalamic region linked to voiding behavior is the lateral hypothalamic area (LHA). Activating glutamatergic LHA neurons results in immediate urinary incontinence (6), whereas activating GABAergic LHA neurons attenuates territorial marking behavior in males (26), (31). However, unlike VMHvl^MC4R^, LHA^MC4R^ neurons failed to label postsynaptic neurons in BAR/LC (Figures S3A and S3B) and, importantly, when stimulated by chemogenetics, failed to affect voiding behaviors (Figure S3C). We conclude that the profound urinary and locomotor phenotypes following chemogenetic stimulation of VMHvl^MC4R^ neurons are mediated by a putative VMHvl-BAR/LC circuit.

**Figure 1.**
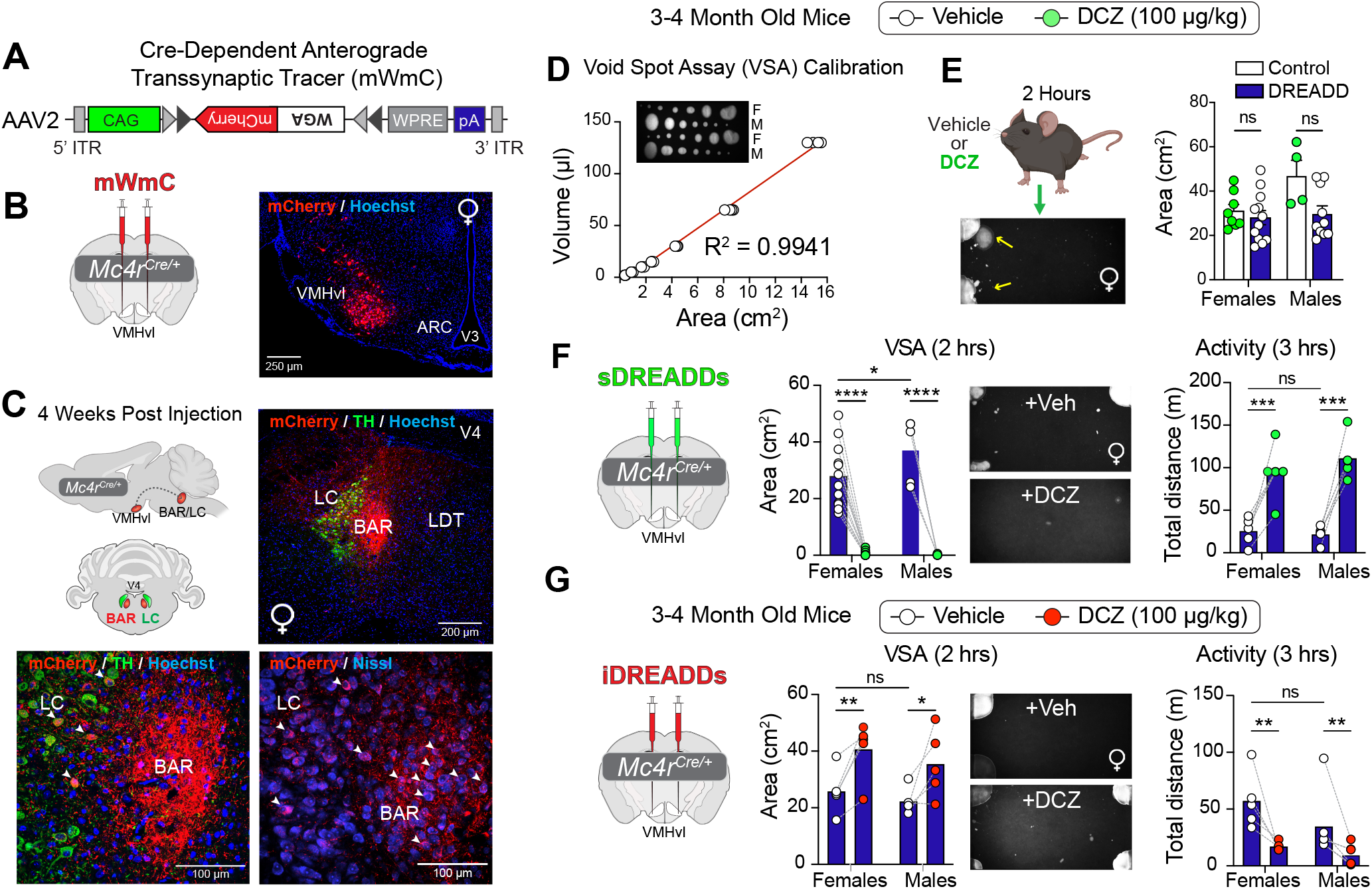
VMHvl-BAR/LC projections are tightly linked with micturition and activity. (A) Schematic showing DIO-mWmC anterograde transsynaptic tracer injected bilaterally into the VMHvl of heterozygous *Mc4r*^*Cre/+*^ 3-4 month old female mice. (B) Representative image showing VMHvl expression of Cre-dependent mWmC tracer, scale bar = 250 µm. (C) Sagittal and coronal schematics of histology results showing strong projections to the BAR/LC hindbrain region. Representative images from a female brain of post-synaptic mCherry staining (white arrowheads) capturing the LC marked by tyrosine hydroxylase (TH), and BAR, marked with Hoechst and Nissl counterstaining; scale bars = 200 µm, upper panels; 100 µm, lower panels. (D) Void spot assay (VSA) calibration using both female and male urine, correlating urine volume with fluorescent area, as described in the Methods (n = 4 replicates from each sex). Non-linear regression test. (E) VSA results and representative images from control mice treated with DCZ (N = 8 females, 4 males) or from *Mc4r*^*Cre/+*^ mice injected with sDREADDs and treated with vehicle (N = 12 females, 10 males); legends above bar graphs. (F) VSA with representative urine image and activity levels with vehicle followed by DCZ treatment (100 µg/kg, IP) measured at least 2 weeks post-injection of sDREADDs (N = 12 females, 5 males). Two-way ANOVA. (G) VSA, with representative urine image and activity levels plus vehicle, followed by DCZ treatment (100 µg/kg, IP), measured at least 2 weeks post-injection of iDREADDs (N = 5 females, 5 males). Two-way ANOVA. *p < 0.05, **p < 0.01, ***p < 0.001, ****p < 0.0001, ns = not significant. Error bars ± SEM. Abbreviations: ARC arcuate nucleus, BAR Barrington nucleus, LC locus coeruleus, LDT lateral dorsal tegmental nucleus, ME5 of mesencephalic trigeminal nucleus, VMHvl ventromedial ventrolateral hypothalamus, V3 third ventricle, V4 fourth ventricle.

### Activating VMHvl^**MC4R**^ Neurons Blocks Urination, Defecation, and Spinal-Brainstem Reflexes

To examine bladder function more fully, in vivo urethane-anesthetized cystometry was performed before and after activation of the VMHvl-BAR/LC circuit under conditions that preserve the spinal-brainstem or spino-bulbo-spinal voiding reflex (32, 33), as described in the Methods. Prior to DCZ treatment, all females exhibited a stereotypic cycle of filling and then voiding during continuous infusion of saline into the bladder. This pattern was immediately and severely altered after activating VMHvl^MC4R^ neurons, leading to an inability to void, a reduction in voiding events, and an elevation in maximal bladder pressure, indicating that when engaged, the VMHvl-BAR/LC circuit blocks spinally-mediated voiding reflexes (Figures 2A and 2B). In vivo uroflow analyses revealed that activating VMHvl^MC4R^ neurons also disrupted defecation as well as voiding, as evidenced by a reduction in fecal events or visible fecal pellets (Figure 2C and Movie S1). The strength of this putative VMHvl-BAR/ LC voiding circuit was readily apparent after challenging mice with a maximal volume of saline prior to the VSA (Figure 2D), after low-dose DCZ treatments (Figures 2E-2G), or after only 7 days post-injection (PI) of sDREADDS into the VMHvl (Figure 2H).

**Figure 2.**
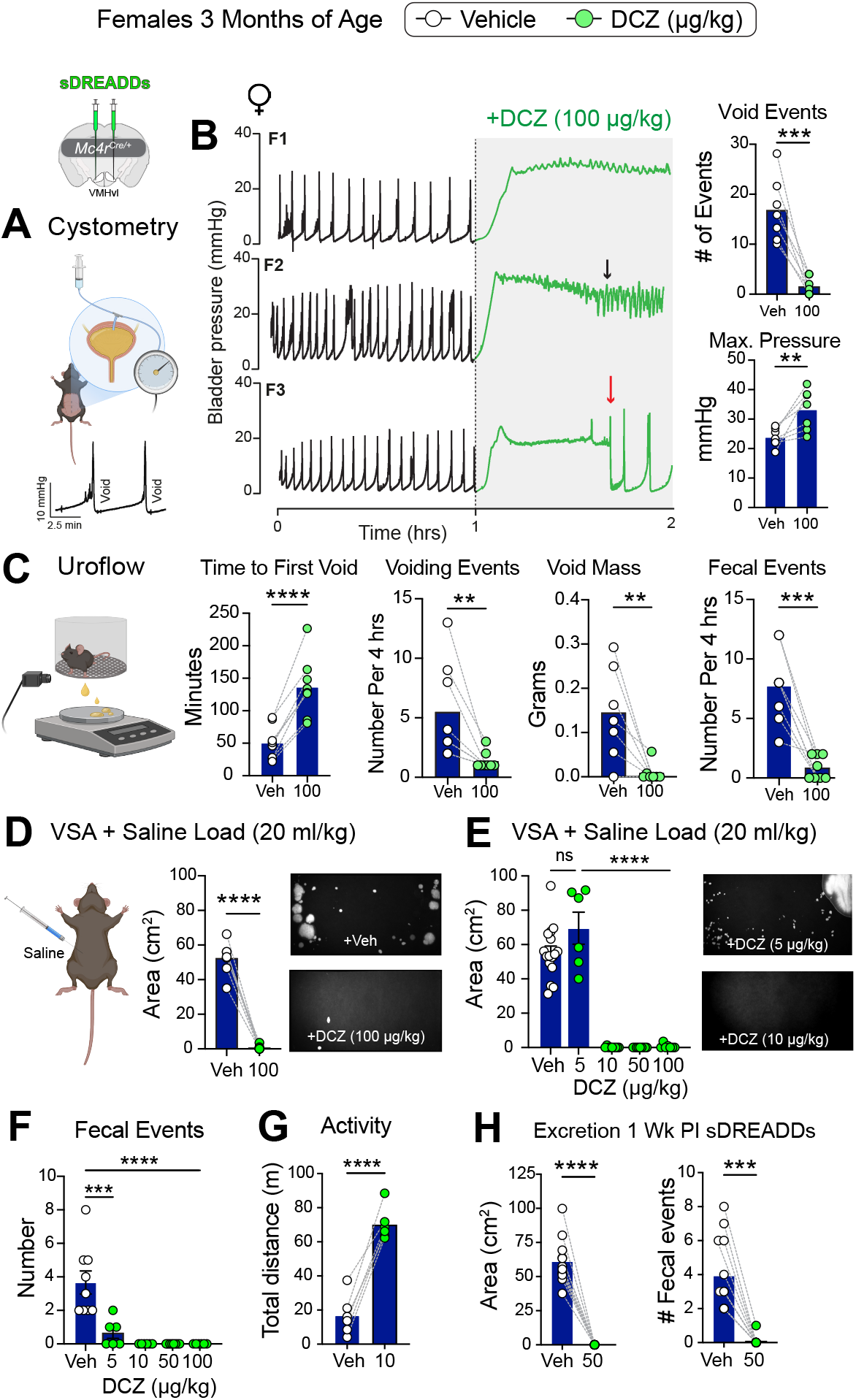
VMHvl^MC4R^ neurons exert a strong central block in bladder reflexes and excretion. (A) Schematic of in vivo anesthetized cystometry and the expected stereotypic pattern of rising pressure followed by voiding. (B) Representative traces before (black traces) and after DCZ treatment (green traces) from three female mice (F1-F3) injected with sDREADDs in the VMHvl. Leakage from the bladder (black arrows) and resumption of the normal voiding pattern (red arrow) are indicated in individual traces. Graphs on the right show the number of void events and maximal pressure averaged from the first 5 voiding events before treatment and then immediately after DCZ treatment (100 µg/ kg; N = 5 females). One-tailed Paired Ratio T-test. (C) Schematic of the uroflow setup and measured parameters over a 4-hour period, including the number of fecal events (N = 7 females). One-tailed, paired Ratio T-test. (D) VSA following a challenge of 20 ml/kg IP saline, as described in methods, with representative images of urine patterns before and after DCZ treatment (100 µg/kg, N = 7 females). One-tailed Paired Ratio T-test. (E) VSA following a saline load with increasing doses of DCZ (5-100 µg/kg, N = 6-16 females), with representative images for 5 and 10 µg/kg DCZ. Note that the data from panel D for vehicle and 100 µg/kg were included in this graph. One-way ANOVA with Tukey’s multiple-comparisons test. (F) Number of fecal events assessed with increasing doses of DCZ (5-100 µg/kg, N = 6-9 females) over a 2-hour period. One-way ANOVA with Tukey’s multiple-comparisons test. (G) Distance traveled following vehicle and 10 µg/kg DCZ (N = 6 females). One-tailed Paired Ratio T-test. (F) VSA, and the number of fecal events in the female cohort following treatment with DCZ (50 µg/kg) just 1 week after injection of sDREADDs into the VMHvl (N = 4 females, 4 males), One-tailed Paired Ratio T-test. **p < 0.01, ***p < 0.001, ****p < 0.0001, ns = not significant. Error bars ± SEM.

Given the marked decrease in defecation following chemogenetic stimulation of VMHvl^MC4R^ neurons, we then asked if colonic motility would similarly be impaired following bead insertion into the distal colon. While expulsion of this exogenous bead is typically complete within 10-15 minutes, activation of VMHvl^MC4R^ neurons by DCZ (100 μg/kg) effectively blocked colonic transit for up to 6 hours (Figure 3A). Even at very low DCZ doses (5 µg/kg), the inability to eliminate the bead in a timely manner persisted (Figure 3A, left graph). Strong inhibition of colonic motility was also observed in aged mice after VMHvl activation (Figure 3A, right graph), which was accompanied by urinary retention and increased locomotion (Figures 3B and S4). The stark delay in colonic transit time, together with the inability to void after bladder filling, suggested that the VMHvl-BAR/ LC projections negatively regulate spinally mediated pelvic organ reflexes. This prompted us to functionally assess the integrity of the colonic spino-bulbo-spinal reflex by measuring visceromotor responses (VMRs) to colorectal distension (CRD), a reflex that persists after decerebration (34). After stimulating VMHvl^MC4R^ neurons, VMR responses were severely blunted even at the highest distension pressures (Figure 3C), mirroring the in vivo cystometry results, which showed a near-complete loss of the voiding reflex despite high bladder pressures. Thus, mice exhibit virtually no visceral responses to high distension pressures in the colon and bladder after activating the VMHvl^MC4R^ node.

**Figure 3.**
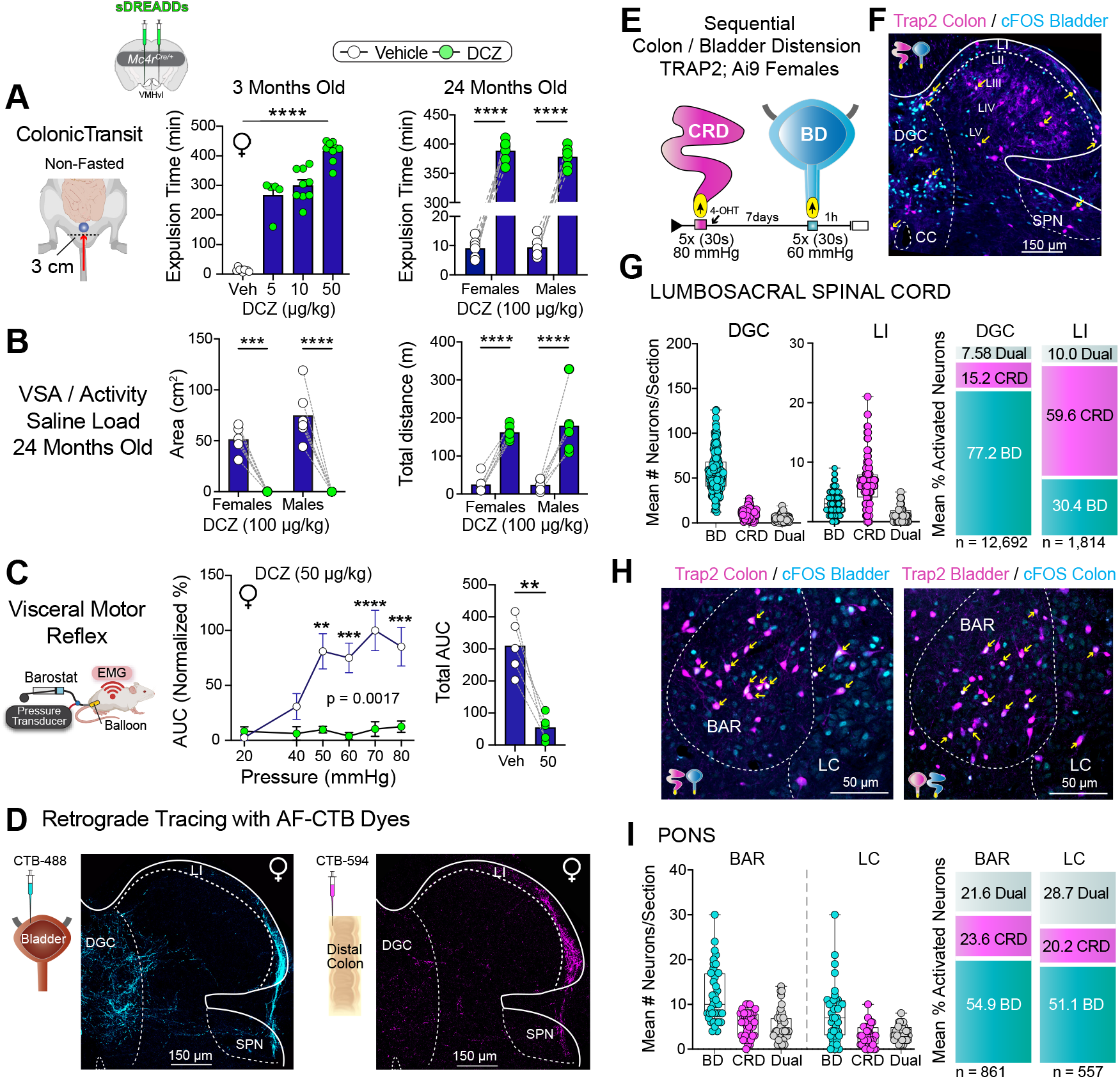
The VMHvl controls pelvic function, with high coactivation of BAR/LC neurons following bladder/colon distensions. (A) Schematic of bead expulsion assay with graph showing colonic transit times in minutes following increasing doses of DCZ in younger female mice, left graph (N = 5 to 9). One-way ANOVA with Tukey’s multiple-comparisons test. Bead expulsion times in aged females and males after DCZ treatment in the right graph (100 µg/kg, N = 6 per group). Two-way ANOVA with Tukey multiple-comparisons test. (B) VSA, and distance traveled in aged cohorts following DCZ treatment (100 µg/kg, N = 6 per group). Two-way ANOVA. (C) Schematic of visceral motor reflex assay to colorectal distension and normalized responses before and after DCZ treatment, with the highest mean level in vehicle-treated females set at 100% (N = 5 females per group). Two-way ANOVA with Tukey multiple-comparisons test. The right graph is the total AUC (normalized) for the untreated and treated cohorts. One-tailed, paired ratio T-test. (D) Retrograde tracing in adult female mice as described in Methods using Alexa Fluor (AF) cholera toxin B (CTB) tracer dyes injected into the bladder (cyan, left panel) or the colorectum (magenta, right panel), with images of the S1 spinal cord shown. Scale bar = 150 µm. (E) Schematic for visualizing activated neurons using Trap2;Ai9 mice (magenta) or cFOS staining (cyan) following sequential distension of the bladder (BD) and colorectum (CRD) at the indicated pressures, performed 7 days apart. (F) Activated neurons in the spinal cord at S1. Scale bar = 100 µm. (G) Number of positive neurons activated by bladder distension (BD, cyan), colonic distension (CRD, magenta), or both (Dual, gray) quantified from spinal cord sections spanning L5-S1, with percentage of each group shown in the right-hand graphs (N=15 female mice). (H) Activated neurons in sections of the BAR/LC region over 1-3 sections. Scale bar = 50 µm. (I) Same description as G for the PONS region with BAR and LC separated (N=16 female mice). **p < 0.01, ***p < 0.001, ****p < 0.0001, ns = not significant. Error bars ± SEM. Abbreviations: BAR Barrington nucleus, CC central cord, DGC dorsal gray commissure, DH dorsal horn, LI lamina layer I of dorsal horn, LC locus coeruleus, SPN sacral parasympathetic nuclei.

That VMHvl neurons exert such a strong effect on both the colon and bladder led us to ask if neurons in the lumbosacral spinal cord and brainstem converge after distension of these two pelvic organs. Initial visualization of bladder and colon afferents using cholera toxin B retrograde tracers (CTB-488 and CTB-594, respectively) revealed discrete regional input into the spinal cord dorsal horn (DH), with dense overlap in lamina I (LI) as well as the dorsal gray commissure (DGC), and within lateral tracts projecting into the SPN (Figure 3D). Activation of spinal and brainstem neurons was quantified following sequential distension of the colorectum (CRD) and bladder (BD). Induction of TRAPed cFOS was detected using *Trap2:Ai9* reporter mice after repeated distensions of one organ. One week later, induction of cFOS was detected by immunostaining after repeated distensions of the alternative organ, as shown by the schematic in Figure 3E. Within the spinal cord, dual-labeled cFOS^+^ neurons – those responding to both colon and bladder distensions – represented a small subset (∼8%) compared to singly-labeled neurons that were more abundant in LI for the colon and in the DGC for the bladder (Figures 3F, 3G and S4A). How-ever, the low convergence of colon- and bladder-activated neurons in the cord nearly tripled in BAR/LC, independent of the method used to visualize cFOS (Figures 3H, 3I and S4B). A similar level of dual-labeled neurons was detected in the PAG (Figure S4C). Thus, the strong convergence of activated neurons in the BAR/LC, together with our findings that chemogenetic activation of the VMHvl essentially blocks all excretion and spinal-brainstem reflexes, suggests that hypothalamic projections to the BAR/LC exert strong central control over adjacent pelvic organs.

### Functional Mapping of the VMHvl-BAR/LC Neural Circuit

To establish that projections from the VMHvl to BAR/LC directly inhibit voiding and promote locomotion, we coupled chemogenetics with the WTR anterograde multi-viral payload system, which involves dual-site injection of three viral vectors (see schematic in Figure 4A and (35)). The tracer vector carrying the modified wheat germ agglutinin (mWGA) and a Tobacco Etch Virus protease (TEVp)-cleavage site (TCS) fused to an inactive payload was injected at the start site – the VMHvl, thereby enabling anterograde transsynaptic delivery of the payload and its activation at the target site – the BAR/LC. There, we delivered the TEVp to the target site, resulting in the cleavage and activation of our delivered payload, the flippase recombinase (FLPo). This sets in motion the final step in this cascade: the expression of FLPo-dependent stimulatory DREADDs also delivered to the target site, thereby enabling chemogenetic manipulation of the BAR/LC neurons innervated by the VMHvl (Figure 4B). For this circuit, chemogenetics rather than optogenetics seemed ideal for monitoring a sustained inhibitory response lasting several hours.

**Fig. 4.**
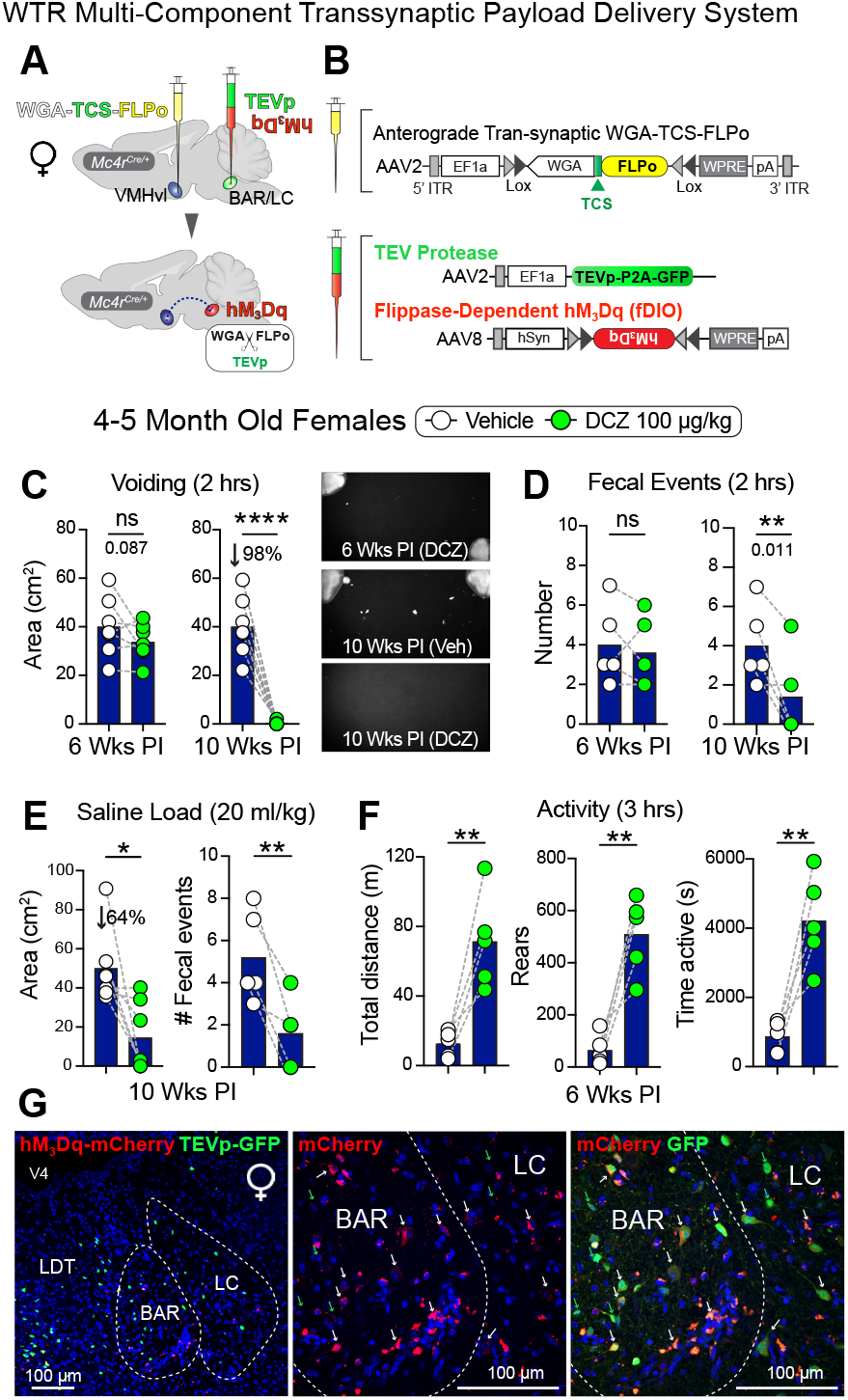
Transsynaptic WTR payload system confirms VMH-vl-BAR/LC as a functional circuit that blocks excretion. (A) Schematic of the three-vector WTR system using functional anterograde transsynaptic tracing to deliver payload from the start site (VMHvl) to the target site (BAR/LC). (B) Components of the three-vector WTR system are depicted, including the anterograde tracer carrying the payload (mWGA-TCS-FLPo) delivered to the VMHvl, and the TEV protease and Flp-dependent sDREADDs viral vectors delivered to BAR/LC or the target site. (C) VSA for 2 hours obtained 6 or 10 weeks post-injection (PI) with representative urine patterns shown at these two time points with vehicle or DCZ (100 µg/kg, N = 6 females). One-tailed Student’s T-test. (D) Number of fecal events in 2 hours quantified at 6-weeks or 10-weeks PI, (N = 5 females). One-tailed Student’s T-test. (E) VSA and number of fecal events quantified with female mice loaded with saline prior to testing at 10 weeks PI (N = 5-7). One-tailed Student’s T-test. (F) Activity parameters shown at 6 weeks PI (N = 5), One-tailed Student’s T-test. (G) Representative images of staining for mCherry (sDREADDs, red) and GFP (TEV protease, green) at low and higher magnifications with overlap indicated by white arrows and shown in the merged image (far right panel). The GFP-only signal is indicated by green arrows. Scale bar = 100 µm. *p < 0.05, **p < 0.01, *****p < 0.0001, ns = not significant. Error bars ± SEM. Abbreviations: BAR Barrington nucleus, LC locus coeruleus, LDT lateral dorsal tegmental nucleus, V4 fourth ventricle.

Weekly monitoring of VSA and physical activity was initiated in a cohort of injected *Mc4r*^*Cre/+*^ females at 5 weeks PI, a time point at which anterograde tracing from the VM-Hvl to BAR/LC can be visualized (see Figures 1B-1D). By 10 weeks PI, voiding was blocked in all females treated with DCZ (Figure 4C). As expected, fecal events were significantly reduced at this time point (Figure 4D); these phenotypes persisted after a saline challenge (Figure 4E). Chemogenetic induction of activity preceded that of blocking excretion, occurring just 6 weeks PI (Figure 4F). After confirming accurate targeting of the WGA-TCS-FLPo tracer vector to the VMHvl (Figure S5A), we found that a substantial number of neurons co-expressed sDREADDs (mCherry signal) and TEVp-GFP (green signal) in the BAR/LC region. By contrast, neurons in the LDT and even the midbrain PAG expressed TEVp-GFP due to substantial viral spreading; however, few or none expressed sDREADDs (Figures 4G and S5B). These functional tracing studies establish that direct projections from VMHvl to BAR/LC are sufficient to motivate activity and block excretion.

### VMHvl-BAR/LC Circuit is Regulated by Inhibitory Neurons and Endogenous Opioids

Next, we wished to determine which neurons in the BAR/ LC region received input from VMHvl neurons. As predicted, nuclear FOS staining in the VMHvl or BAR/LC was observed only after DCZ treatment in female mice injected with sDREADDs, but not after iDREADD injection or vehicle treatment (Figure 5A and Figure S6A). Histological analyses revealed that the majority of *cFos*^+^ neurons in BAR expressed *Vgat* and *Lhx1*, two markers of inhibitory neurons (Figure 5B, C, and Figure S6B). To test whether inhibitory neurons in BAR/LC are sufficient to block urination and defecation following stimulation of VMHvl neurons, *Vgat*^+^ neurons were selectively ablated using a Cre-dependent diphtheria toxin A (DTA) viral vector delivered bilaterally to the BAR/LC in *Vgat*^*Cre/+*^*;GFP* mice that also received sDRE-ADDs into the VMHvl (Figure 5D). Reducing the number of BAR/LC inhibitory neurons had no effect on baseline voiding (vehicle-treated) but restored near-normal patterns in both sexes after DCZ treatment at low doses. However, at a higher DCZ dose, this breakthrough was observed only in male mice (Figure 5E). The number of fecal events and colonic transit time, as measured by bead expulsion, showed a similar but less dramatic sex-specific breakthrough (Figure 5F, G). By contrast, the increased activity following DCZ treatment persisted and was equivalent in all groups (Figure 5H). Thus, *Vgat*^*+*^ inhibitory neurons in BAR/LC are an obligatory component of this hypothalamic-hindbrain voiding circuit.

**Figure 5.**
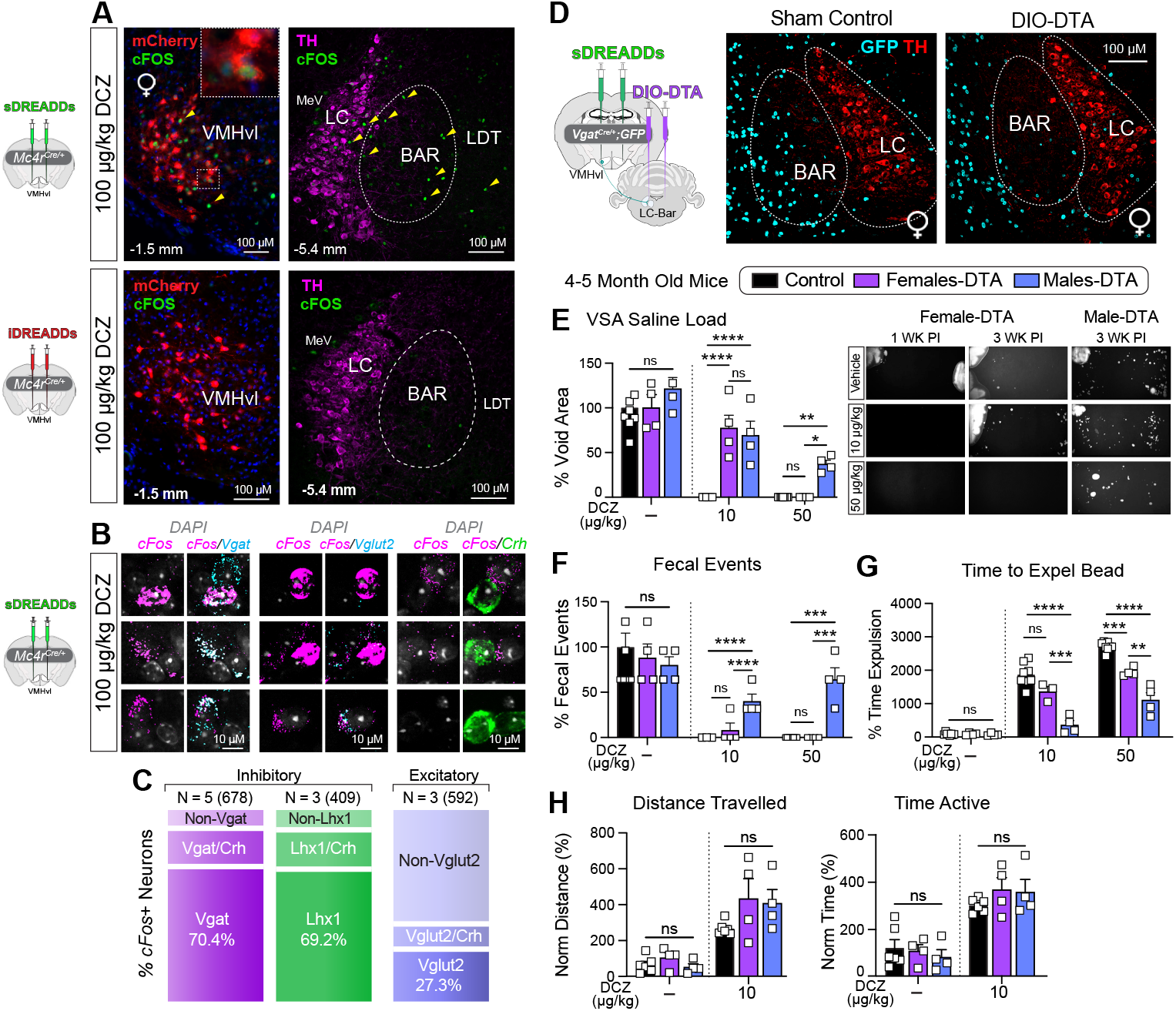
Ablating inhibitory neurons in the BAR/LC partially reverses the excretion block imposed by VMHvl neurons. (A) Representative images of nuclear staining of cFOS-positive neurons (green, yellow arrowheads) in the VMHvl injection site after stimulating (upper panels) or inhibiting (lower panels) VMHvl^MC4R^ neurons with DCZ (100 µg/kg), as marked by mCherry staining. Tyrosine hydroxylase (TH, magenta) staining marks the LC region. The magnified area in the upper left-hand panel shows nuclear cFOS staining. Scale bars = 100 µm. (B) Representative images in BAR of overlapping *cFos* with *Vgat, Vglut2*, or *Crh*, after stimulation of VMHvl^MC4R^ neurons with DCZ (100 µg/kg). Scale bar = 10 µm. (C) Stacked bar graphs indicating percentage of overlapping *cFos*+ neurons with biological replicates (N) and neurons counted (n) indicated above each bar. (D) Schematic of sDREADDs into VMHvl and DIO-DTA viral infections into BAR/LC region in *VgatCre/+;GFP* reporter mice. Representative images of female mice 4 weeks PI of sDREADDs and DIO-DTA, stained for VGAT+ neurons (GFP, cyan) and tyrosine hydroxylase (TH, red) marking the LC. Scale bar = 100 µm. (E) VSA (2 hrs) following saline load in male and female mice assayed 3 weeks PI of viral vectors, after treatment with vehicle (-), 10 or 50 µg/kg of DCZ (N = 4 females and 4 males). All values were normalized to the vehicle-treated sham cohort, with the mean value set to 100% (N = 8). One-way ANOVA with Šidák multiple comparisons for the three DCZ-treated groups, including sham, DTA-females, and DTA-males. Representative images to the right show urine patterns for 1 and 3 Weeks PI. (F) Same as panel E, % of fecal events. (G) Same as panel E, % time of colonic bead expulsion. (H) Same as panel E, % distance traveled and time active. *p < 0.05, **p < 0.01, ***p<0.001, ****p < 0.0001, ns = not significant. Error bars ± SEM. Abbreviations: BAR Barrington nucleus, LC locus coeruleus, LDT lateral dorsal tegmental nucleus, ME5 of mesencephalic trigeminal nucleus, VMHvl ventromedial ventrolateral hypothalamus, V4 fourth ventricle.

We then asked whether pharmacological intervention of the autonomic nervous system would similarly reverse the strong voiding phenotypes. Prior to this, catecholamines and other HPA stress hormones were measured following DCZ treatment. Somewhat unexpectedly, the large chemogenic-induced changes in movement and excretion resulted in only a modest change in one of two catecholamines (norepinephrine), but a significant, substantial increase in prolactin (Figure 6A), a pituitary hormone that rises during the pre-ovulatory surge in rats (36) and mice (37), when mate-seeking behavior is triggered. Somewhat surprisingly, blocking peripheral sympathetic tone with guanethidine (GUA) (38) had little effect on the VMHvl-BAR/LC voiding circuit (Figure 6B), consistent with the fact that ACTH and catecholamines, two humoral outputs of sympathetic activation, are largely unchanged. Furthermore, bethanechol, a cholinergic drug that targets parasympathetic bladder innervation and is used to promote urination, failed to counteract the effects of VMHvl activation on urination but did promote uncontrolled fecal emptying or diarrhea, as shown by others (Figure 6C).

**Figure 6.**
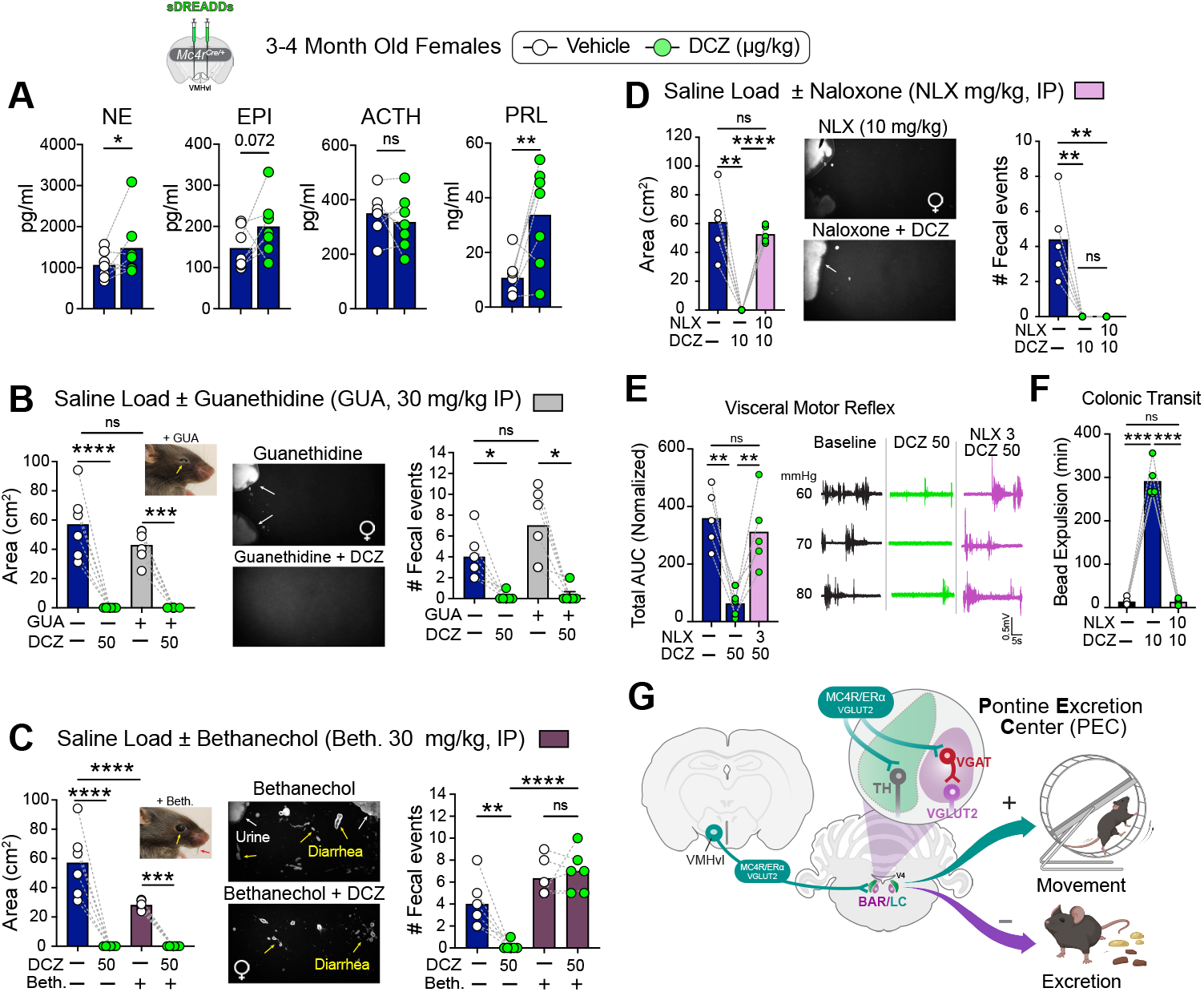
Blocking opioid signaling restores bladder and colonic sensations opposing the VMHvl^MC4R^-BAR/LC circuit. (A) Levels of catecholamines and pituitary hormones before and after DCZ treatment (N = 7 females). One-tailed Paired Ratio T-test. (B) VSA (left graph) and number of fecal events (right graph) in females treated with guanethidine (GUA, 30 mg/kg, IP, grey bars) 2 hours prior to activation of VMHvl^MC4R^ neurons with DCZ (100 µg/kg). GUA-induced pupil constriction (yellow arrow) is shown in the insert for one mouse. Representative images in the middle panels show urine patterns with GUA alone or with DCZ, N = 6 females. Repeated One-Way ANOVA with Tukey’s multiple comparisons. (C) VSA (far left panel), and number of fecal events (far right panel) in females treated with bethanechol (Beth, 30 mg/kg, IP, purple bars) just prior to activation of VMHvl^MC4R^ neurons with DCZ (100 µg/kg). Bethanechol-induced pupil dilation (yellow arrow) and saliva production (red arrow) are shown in the insert for one mouse. The representative images in the middle panels show urine patterns and diarrhea marks (yellow arrows) after bethanechol alone or with DCZ, N = 6 females. RM One-Way ANOVA with Tukey’s multiple comparisons. (D) VSA (far left panel) and number of fecal events (far right panel) in females treated with naloxone (NLX, 10 mg/kg, IP, pink bars) 10 minutes prior to activation of VMHvl^MC4R^ neurons with DCZ (10 µg/kg), N = 5 females. Representative images in the middle panels show urine pattern marks naloxone alone or with DCZ. RM One-Way ANOVA with Tukey’s multiple comparisons. (E) Total AUC for VMR responses to increasing distension pressures, normalized to vehicle-treated group, with representative traces of EMG recording at 60, 70, and 80 mmHg shown for one animal. N = 5 females. (F) Time for colonic bead expulsion in females treated with vehicle or naloxone (NLX, 10 mg/kg, IP, pink bars) 30 minutes prior to activation of VMHvl^MC4R^ neurons with DCZ (10 µg/kg), N = 5 females. RM One-Way ANOVA with Holm-Šidák multiple comparisons. (G) Schematic of VMHvl input into the hindbrain Pontine Excretion Center (PEC). *p < 0.05, **p < 0.01, ***p<0.001, ****p < 0.0001, ns = not significant. Error bars ± SEM.

The inability of these pharmacological manipulations to reverse the effects of DCZ on voiding suggests that activation of this VMHvl-BAR/LC circuit engages pathways beyond the descending autonomic system. Moreover, the loss of normal spinal-brainstem reflexes to high colonic and bladder distension pressures in our VMR and cystometry studies suggested that a hypo-nociceptive state is achieved following activation of VMHvl^MC4R^ neurons and that normal reflexes to painful or noxious stimuli are effectively muted. Involvement of endogenous opioid pathways was tested by a single injection of the pan-opioid antagonist naloxone (Narcan), administered prior to DCZ treatment. Naloxone completely reversed the effects of DCZ, restoring normal urination in saline-loaded females without affecting locomotor activity (Figure 6D and Movie S1). Similarly, naloxone reversed DCZ-mediated suppression of VMR responses following colorectal distensions at noxious pressures (60-80 mmHg) (Figures 6E and S7). As observed for micturition, naloxone-induced reversal of DCZ was equally dramatic for colonic transit, decreasing bead expulsion time from hours to minutes; fecal events remained suppressed (Figures 6D and 6F). Together, these pharmacological data infer that activating VMHvl^MC4R^ neurons triggers endogenous endorphin signaling, thus dampening visceral sensations and spinal-brainstem reflexes in the bladder and colon.

## DISCUSSION

Our findings provide new insights into how a distinct hormone-responsive hypothalamic node coordinates a suite of voluntary behaviors, prioritizing exploratory-like behavior while suppressing the time-consuming, spatially restricted choice to excrete (Figure 6G). The ability to essentially block all excretion in young and aged animals after activation of the VMHvl^MC4R^-BAR/LC circuit, even at very low DCZ doses, underscores the strength of this circuit as mapped by our functional tracing studies. Further, this block persists when challenged with increased volume load or colonic bead insertion, and, importantly, under noxious conditions that would normally promote acute colonic and bladder visceral motor reflexes and pelvic discomfort.

That the VMHvl-BAR/LC circuit exerts such a strong effect on both urination and defecation builds on prior studies implicating this brainstem region in coordinating these two pelvic functions. Indeed, crosstalk between the BAR/LC region and the convergence of pelvic visceral afferents was suggested over 20 years ago (40), and may underlie the high, shared incidence of irritable bowel syndrome (IBS) and bladder pain, particularly in women (41, 42). Recent studies by Grundy et al. show that TNBS chemical induction of colonic hypersensitivity leads to hypersensitivity of bladder afferent pathways and abnormal voiding (43). Conversely, Kim et al. demonstrate that bladder pain induced by intravesical administration of the LL-37 antimicrobial peptide resulted in colonic hypersensitivity (44). Bifurcating or dichotomizing spinal afferents between the lumbosacral (LS) dorsal root ganglia (DRG) and the bladder and colon were originally proposed to mediate pelvic organ cross-sensitization (45), an observation confirmed by subsequent studies showing 10-15% convergence in the lumbosacral DRG afferents from colon and bladder (43, 44). Interestingly, we also observe a minor population of spinal cord neurons that are dual-labeled and coactivated following sequential distensions of the bladder and colon. However, in the BAR/LC region, the percentage of convergent neurons increases to well over 20%, with nearly equivalent percentages detected in either the BAR or the LC. Thus, the BAR/LC likely serves as a central gatekeeper for crosstalk between the bladder and colon. When coupled with our functional data, we posit that the BAR/LC can no longer be viewed narrowly as a pontine micturition-only center (PMC) or a defecation-only brain area (DBA) (12), but should be broadened and redefined as the pontine excretion center (PEC, Figure 6G). Whether the BAR/LC regulates other organs innervated by the pelvic splanchnic nerves remains intriguing, given the high comorbidity of bowel and bladder symptoms among women with other forms of pelvic pain, such as endometriosis (46). If true, the functional definition of the BAR/LC will continue to evolve.

Our study now identifies the VMHvl as an important hypothalamic region in the central control of excretion, significantly expanding the repertoire of functions for this hormone-responsive node. Other regions implicated in urination include the LHA and the medial preoptic area (MPOA) (7, 31), both of which appear particularly relevant to male scent marking (26). The same is not true for the VMHvl. The connection between excitatory neurons in the VMHvl^MC4R^ and the BAR/LC is consistent with recent work showing that, in addition to strong projections from the LHA and midbrain vlPAG to BAR, sparse yet clear retrograde labeling from either the BAR (5, 6) or the colon (12) targets the VMHvl. We suggest that the small number of scattered *Vgat*^+^ inhibitory neurons in the BAR/LC region, acting locally rather than on the spinal cord, are positioned to modulate descending (5, 12) or ascending (11) projections to and from these two pelvic organs, a proposal supported by an older observation that pharmacological activation of inhibitory neurons in the BAR/LC region increases urinary retention (47).

Chemogenetics proved ideal for parsing out three distinct volitional endpoints – movement, urination and defecation. Indeed, DCZ dosing revealed that excretion was restored in males but not in females after ablation of BAR/LC^VGAT^ neurons, suggesting that either the quantity or the intrinsic properties of VMHvl^MC4R^ neurons differ by sex. Further, the ability to uncouple activity from excretion after BAR/ LC^VGAT^ ablation or naloxone treatment also suggests that an excitatory neuronal cluster shared between BAR and LC, such as the *Penk*^*+*^*/Mc4r*^*+*^ cluster (4), promotes movement. Whether the TH^+^ neurons in the LC targeted by the VMHvl^MC4R^ or second-order projections from BAR/LC to other brain regions initiate movement remains to be determined. Nonetheless, our study highlights the utility of chemogenetics for examining intrinsic robust physiological responses that span long timescales, lasting hours rather than minutes or seconds, an aspect that is more difficult to achieve with optogenetics.

For reproductively active females, we suggest that the transient coordination of exploratory movement and excretion seems perfectly designed to facilitate mate-seeking, especially during a shortened or seasonally restricted breeding period. We would predict that the pre-ovulatory surge in estrogen activates the VMHvl-BAR/LC circuit to promote movement while reducing pelvic pain, in contrast to its action, specifically in the distal colon, where estrogen amplifies visceral gut sensitivity (48). This implies that, in addition to estrogen, other humoral inputs likely activate the VMHvl-BAR/LC circuit; identifying them will be important for the future translation of our findings to address urinary and fecal incontinence. It will also be of interest to know if improvement of lower urinary tract symptoms with hormone replacement therapy in older estrogen-depleted females as bladder control degrades (49), involves central mechanisms in addition to local effects. Regardless, restoring colonic sensation, voiding, and colonic propulsion with a single dose of naloxone following DCZ treatment is striking, suggesting that endogenous opioids are directly linked to the phenotypes observed following VMHvl^MC4R^ activation. In this regard, the decades-old observation that prolactin levels rise significantly after β-endorphin treatment (50) is highly relevant, given that we also observe a rise in circulating prolactin to similar levels following activation of VMHvl^MC4R^ neurons. Further work will be needed to pinpoint the source and subsequent actions of endogenous endorphins on this neuroendocrine-hindbrain circuit.

## METHODS

### Ethics

All experiments were approved and conducted in accordance with the guidelines set by the Institutional Animal Care Committees (IACUC) at University of California, San Francisco (UCSF), the University of Wisconsin–Madison, or the South Australian Health and Medical Research Institute (SAHMRI), the National Institutes of Health Guide for Care and Use of Laboratory Animals, and the recommendations of the International Association for the Study of Pain. Mice were housed on a 12-hour light/dark cycle with ad libitum access to food and water unless otherwise noted.

### Mice

*Mc4r-t2a-Cre* mice (51) were a gift from B. Lowell and were maintained in the laboratory at UCSF on a C57BL/6J background. *Slc32a1*(*Vgat)-ires-Cre* knock-in mice (52) (gift from K. Yackle and E. Feinberg) were crossed to *CAG-LSL-Sun1/sfGFP* mice (53) (gift from K. Yackle) and maintained on a mixed background. Heterozygous mice were used in all experiments involving Cre knock-in lines. 22-24-month-old aged C57BL/6 mice were obtained through the National Institute on Aging (NIA) Aged Rodent Colony Program, available to NIA-funded projects.

Female TRAP2/tdTomato mice (10-14 weeks of age) were used for the colon and bladder co-activation assay. A colony of TRAP2 mice was established at SAHMRI via rederivation from TRAP2 mice provided by Irina Vetter at the University of Queensland (originally sourced from Jackson Laboratory, Strain #: 030323) (54). TRAP2 mice were crossed to Ai9 mice (SAHMRI colony rederived from Jackson Laboratory, Strain #: 007909) that express tdTomato under Cre control.

### Stereotaxic Surgeries

Adult mice (8-16 weeks old) were secured under isoflurane anesthesia in a Model 1900 stereotaxic frame (David Kopff Instruments), and viral vectors were injected bilaterally at the following coordinates. For the VMHvl, anterior–posterior: bregma −1.48 mm, mediolateral: bregma ±0.85 mm, dorsoventral: bregma −6 mm. For the BAR, anterior–posterior: bregma −5.2 mm, mediolateral: bregma ±0.7 mm, dorsoventral: bregma −4.1 mm. For the LHA, anteri-or–posterior: bregma −1.48 mm, mediolateral: bregma ±0.85 mm, dorsoventral: bregma −5.2 mm.

For Cre-dependent anterograde transsynaptic tracing, mice received bilateral injections of AAV2-CAG-DIO-mWGA-mCherry-WPRE (mWmC, 150 nl, Virovek Inc, Cat # 117-UCSF86), as previously described by X.D. (27). A codon-optimized wheat germ agglutinin (mWGA) sequence was synthesized commercially (GENEWIZ Inc., South Plainfield, NJ), fused with mCherry, and then cloned into an AAV-CAG-overexpression-WPRE vector. For the transsynaptic payload delivery system, the WTR toolkit was employed as previously detailed in (35). Briefly, the AAV2-EF1α-DIO-mWGA-TEVcs-Flpo (titer: 5.7×1012 vg/ml, Packgene) was generated by fusing in-frame mWGA with a Tobacco Etch Virus protease (TEVp) cleavage site (TEVcs) recognition sequence (GAGAACCT-GTACTTCCAGGGC) and FLPo recombinase. The TEVp coding sequence was amplified from pAAV-flex-taCasp3-TEVp (Addgene #45580) and then subcloned to produce AAV2-EF1α-TEVp-P2A-GFP (titer: 2×1012 vg/ml, Packgene). Mice received bilateral injections of AAV2-EF1α-DIO-mWGA-TEVcs-Flpo into the VMHvl (300 nl), and AAV2-EF1α-TEVp-P2A-GFP (140 nl) and AAV8-hSyn-fDIO-hM3D(Gq)-mCherry-WPREpA (Addgene #154868, 60 nl) into the BAR.

For chemogenetic experiments, mice were bilaterally injected with AAV2-hSyn-DIO-hM3D(Gq)-mCherry (Addgene #44361, 110 nl), AAV2-hSyn-hM3D(Gq)-mCherry (Addgene #50474, 100 nl), or AAV2-hSyn-DIO-hM4D(Gi)-mCherry (Addgene #44362, 250 nl) DREADD viral vectors. For experiments combining ablation of GABAergic neurons in BAR with chemogenetic activation of the VMHvl, mice received bilateral injections of AAV2-hSyn-hM-3D(Gq)-mCherry (Addgene #50474, 100 nl) into the VMHvl and AAVDJ-CAG-DIO-DTA-WPRE-hGH (BrainVTA #PT-0775, 80 nl) or AAV2-hSyn-DIO-mCherry (Addgene #50459, 80 nl) into the BAR. The reagents used in this study are listed in Table S1.

For all surgeries, mice were allowed to recover for at least 1 week before any behavioral assays. For transsynaptic labeling, mice were allowed to express the reporter for 4-5 weeks before tissue collection. At the conclusion of all in vivo experiments, mice were perfused with PBS followed by 4% PFA, and the brains were cryo-preserved to confirm proper targeting. Any mice without correctly targeted fluorescent protein expression were excluded from subsequent analyses.

### Void Spot Assay

Void spot assay (VSA) was conducted to study mouse micturition behavior noninvasively (29). Testing cages were prepared immediately prior to each test using clean, standard mouse cages without bedding and lined with a fresh, dry sheet of filter paper (Cytiva Whatman, #1001-917). Urine from each void is absorbed by the paper, enabling subsequent quantification of the resulting urine spot pattern. A small paper ball was placed on the cage floor to occupy the mouse during the test; food and water were withheld to prevent contamination and eliminate variation in bladder filling and voiding resulting from differences in water intake among individuals. Mice were allowed to acclimate to the behavior testing room in their own cages for 30 min prior to being treated and introduced to the paper-lined cages for the 2-hour test. When saline ‘volume-load’ was performed, mice were administered an intraperitoneal (IP) injection of saline at 20 ml/kg (55, 56) and returned to their home cages for 15 min before being tested to avoid contaminating the experimental arena (filter paper) with fluid drips from the injection or small stress-induced voids. VSAs were always run at the same time of day, during the first phase of the light-on cycle (resting phase). For chemogenetic experiments, mice were injected IP with either saline (vehicle) or Deschloroclozapine dihydrochloride (DCZ; Hello Bio #HB9126) immediately before starting the VSA. A DCZ working solution (0.04 mg/ml) was prepared daily from a 0.5 mg/ml DCZ stock and administered to mice at a dose of 5, 10, 50, or 100 μg/kg.

Each mouse underwent VSA testing following control (IP saline) and chemogenetic (IP DCZ) treatments, counterbalanced across two separate days, with DCZ administered on either the first or second day of testing. For several conditions, VSAs were repeated and measurements taken weekly over several weeks, as was the case for testing the WTR Payload system (beginning at post-injection (PI) Week 5) and the DTA ablation studies (beginning at PI Week 1). Papers were allowed to dry, and urine spots were illuminated and imaged under UV light (ChemiDoc MP Imaging System, Bio-Rad). Using ImageJ software (NIH), images were scaled, thresholded, and analyzed to determine the number of urine spots and the area covered by urine. Urine volume was calculated from urine-covered area (pixel area detected) using second-order polynomial coefficients determined on a calibration curve. Calibration data consisted of 4 replicates of 7 different known urine volumes on filter paper (2 replicates using pooled female urine and 2 replicates using pooled male urine; Figure 1E). In addition, the number of fecal events (fecal pellets) was also counted at the end of each 2-hour test.

### Uroflow Analyses

Uroflowmetry was performed at the University of Wisconsin-Madison as described previously (57). Briefly, mice were housed in the same room as the testing. Mice underwent uroflowmetry testing on two consecutive days: one day treated with an IP injection of saline and one day treated with 100 μg/kg DCZ. Treatments were counterbalanced so that DCZ was administered on either the first or second day of testing. Following treatment, mice were immediately placed in uroflowmetry chambers equipped with Raspberry Pi units interfaced to a balance, enabling video capture of each void along with void mass using Void Sorcerer open-access design and software (58). Mice were tested over a 4-hour period with access to water but not food. All testing was conducted during the same time of day. Uroflowmetry data were analyzed by an individual blinded to treatment conditions. Only urine events that fell without inference from the grid floor bars or without co-occurrence of a fecal event were used in all analyses. Urine mass was converted to urine volume by dividing the change in urine mass by 1.0046 g/ ml. Flow rate was then calculated as the change in volume over the change in time for urine events.

### In Vivo Cystometry

Anesthetized cystometry was performed as previously described (57). Briefly, mice were anesthetized with a subcutaneous injection of urethane dissolved in saline (AC32554-0500, Fisher, Waltham, MA) at a dose of 1.43 g urethane/kg body weight, prepared from a fresh stock of 86 mg/ml in saline. When unresponsive to toe pinch (at least 30 minutes), the abdomen was opened, and a purse-string suture was placed in the dome of the bladder using a curved needle and 6–0 Silk (501180809, Fisher). A 25 G 1.5-inch needle was fitted with a catheter of PE-50 tubing (NC9140178, Fisher) cut to the length of the needle and flared at the bottom. The needle and catheter were inserted into the dome of the bladder. The needle was removed, and the purse string suture was tied to secure the catheter in place. The abdominal wall and skin were closed with a suture, and the mouse was placed on a heating pad for ∼60 min. Mice were then connected to an in-line pressure transducer and infusion pump. Saline was infused into the bladder at a rate of 0.8 ml/hour, and intravesical pressure was recorded using an MLT844 physiological pressure transducer (ADInstruments, Colorado Springs, CO) connected to an FE221 Bridge Amp (ADInstruments) with a Power Lab 2/26 (PL2602) data acquisition system. Cystometrograms were recorded and analyzed using LabChart software (ADInstruments). Recordings were conducted for ∼1 hour to achieve a steady pattern. Mice received an IP injection of saline control prior to the start of cystometry for the first 1-hour recording and then received an IP injection of DCZ at 100 µg/kg for the second 1-hour recording. DCZ working solutions were prepared fresh daily from a 0.5 mg/ml DCZ stock. 5 consecutive voids were analyzed and averaged per animal by an individual blinded to treatment conditions. Parameters measured from cystometrograms included the number of void events during the 1-hour testing period and maximum bladder pressure before and after DCZ. If intravesical pressure did not build over the hour and was not recovered with a subsequent repositioning of the catheter or if the animal failed to elicit voids, then the mouse was excluded from analysis. After cystometry, mice were euthanized with CO2, perfused with PBS, then 4% PFA, and necropsy was immediately performed to collect brain tissue.

### Colonic Transit Time Assay

Mice were fed ad libitum and habituated to clean cages with material (Kimwipes) for 1 hour. Following saline or DCZ treatment (IP) at doses of 5, 10, 50, and 100 μg/kg, mice were briefly anesthetized with isoflurane. The colonic contents were thoroughly flushed out with a 100 µl saline infusion using a lightly lubricated, flexible, blunt-tipped feeding tube. A 3 mm glass bead (Z143928, Sigma) was then gently inserted into the colon to a depth of 3 cm as determined by using a marked fistula. The time from bead insertion to expulsion, as determined visually, was recorded. Saline-treated mice were monitored every 5 min for bead expulsion, and DCZ-treated mice were monitored every 5 min for the first 20 min, then every 15 min thereafter.

### In Vivo Assessment of Colonic Pain

Visceromotor responses (VMR) to colorectal distension (CRD) were assessed using electromyography (EMG) in fully awake animals, as previously described (48). To allow repeated measurements over time in the same animal, a wireless transmitter (ETA-F10; Data Sciences International, New Brighton, MN, USA) was placed subcutaneously, with the leads inserted into the abdominal musculature. Animals were housed individually post-surgery and allowed to recover for at least 10 days before VMR tests. Under brief anesthesia, the distal colon was cleared of its contents, and a lubricated 2 cm long balloon was gently inserted into the colorectum up to 0.25 cm beyond the anal verge. Once the balloon catheter was secured to the base of the tail, the animal was moved to a restrainer and allowed to recover from anesthesia for 15 minutes. The balloon catheter was then connected to a barostat (Isobar 3, G&J Electronics, Willowdale, Canada) for graded, pressure-controlled delivery of the balloon-distension sequence. The distensions were 20-sec long with 2-min intervals and applied in the following sequence: 20, 40, 50, 60, 70, and 80 mmHg. Animals were treated with saline or DCZ (50 µg/kg) for 15 minutes before the start of the distension sequence. EMG recordings in response to the distensions were relayed to Ponemah Software (Data Sciences International) data acquisition system and analyzed offline using SpikEB Software (CED). The magnitude of VMR responses to each distension pressure was quantified by computing the area under the curve (AUC) during distension (20 seconds), corrected for baseline EMG activity from the 20 seconds immediately before distension. All VMR values were normalized to the average maximum distension values at baseline (without DCZ) set to 100%, and absolute zero was set to 0%. Total AUC for each animal was calculated as the sum of the normalized VMR responses at distension pressures of 50, 60, 70, and 80 mmHg. Repeated VMR assays in the same animal were spaced at least 5 days apart.

### Activity Monitoring

Ambulatory activity in mice was recorded over a 3-hour period and quantified using the ANY-maze behavioral tracking system (Stoelting, v.7.65) as previously described (20). Before all measurements, mice were acclimatized to the ANY-maze chambers for at least 1 hour. Recordings were always conducted at the same time of day, during the first phase of the light-on cycle (resting phase).

### Circulating Pituitary Hormone/Catecholamine Measurements

*Mc4r*^*Cre/+*^ female mice injected with Cre-dependent sDREADDs into the VMHvl were treated with 100 μg/kg DCZ or vehicle (IP) 15 min before blood collection. Animals were briefly anesthetized with 1.5% isoflurane, and submandibular venous blood was collected in EDTA-coated tubes (Microvette CB 300) between ZT 2–5 and immediately treated with EGTA–glutathione for catecholamine analysis or aprotinin (500–1,000 KIU/ml blood) for pituitary hormone analysis. Plasma was isolated by centrifugation at 1,500 rcf for 8 minutes at 4°C and stored at −80°C. Circulating catecholamines and pituitary hormones were measured by the VUMC Hormone Assay and Analytic Services Core. Catecholamine levels were measured by high-performance liquid chromatography, and serum levels of pituitary hormones were measured by multiplex fluorescent Luminex Assay, as previously described (59).

### Pharmacological Treatments

The peripheral adrenergic neuron blocker, guanethidine monosulfate (Sigma-Aldrich, 1301801; 30 mg/kg, IP) was injected 2 hours before the saline load (20 ml/kg) and VSA. The parasympathomimetic muscarinic agonist bethanechol (Sigma-Aldrich, 1071009) was administered IP at 30 mg/kg immediately before the saline load and VSA. In both cases, pharmacological treatments were administered concomitantly with treatment for chemogenetic activation (vehicle vs. DCZ). The opioid antagonist naloxone (Sigma-Aldrich, N7758) was injected IP at either 3 mg/kg for 10 min before DCZ treatment in the VMR assays or at 10 mg/kg 15 min after the saline load and 10 min before DCZ in the VSAs.

### Colon And Bladder Co-Activation Assay

Female TRAP2/tdTomato mice were maintained under isoflurane anesthesia during *in vivo* colorectal distension (CRD) or bladder distension (BD). For CRD (11, 60), the balloon was inflated to 80 mmHg using a syringe attached to a three-way tap and monitored with a sphygmomanometer. Distension was held for 30 seconds, then released for 5 seconds. This sequence was repeated 5 times. For BD (61), a catheter (PE 50 tubing) was inserted into the bladder via the urethra. A fine silk suture was loosely tied around the catheter and the urethral opening to secure the tube and limit pressure loss from leakage. The catheter was attached to an empty 5 ml syringe and sphygmomanometer via a three-way tap. The bladder was inflated to 60mmHg for 30 seconds, then released for 5 seconds. This sequence was repeated 5 times.

4-hydroxytamoxifen (4-OHT, Sigma-Aldrich) was dissolved in 100% ethanol to a final concentration of 25 mg/ml by vortexing at 37°C for 15-20 min and stored at -20°C. For use on the day of injection, the dissolved 4-OHT was combined with Chen oil (a mixture of 4 parts sunflower seed oil and 1 part castor oil) at a concentration of 10 mg/ml and vortexed for 15 min. The ethanol was then evaporated using nitrogen evaporation at 37°C for 10min. The final 10 mg/ml 4-OHT solution was then IP injected at a dose of 50 mg/kg immediately after the final colorectal or bladder distension. Mice were individually housed and monitored daily for any adverse responses to organ distension or 4-OHT injection. Mice were given 1 week to allow for tdTomato expression before undergoing *in vivo* distension of the alternate organ. A separate group of mice received a 4-OHT injection without organ distension and without stimulation 1 week later. Afterward, mice were returned to their home cage for 1 hour before undergoing transcardial perfusion.

### Retrograde Tracing from the Colon and Bladder

Retrograde tracing using cholera toxin subunit B (CTB) was performed to identify the spinal sensory afferent input from the colorectum and bladder to the lumbosacral spinal cord and to correlate the distribution of afferent projections with the distribution of neurons activated by colorectal or bladder stimulation. A separate group of mice underwent CTB-retrograde tracing from the colorectum and bladder 1 week after *in vivo* colon or bladder distension and 4-OHT injection. Retrograde tracing using CTB (0.5% in 0.1 M phosphate buffer) conjugated to Alexa Fluor 488 or 647 (CTB-488: Invitrogen, C22841; CTB-647: Invitrogen, C34778) was performed from the colorectal and bladder walls as previously described (43, 60, 61). Injections into the colorectal wall were made at 2 sites, covering 0.5 cm distal and 1.5 cm proximal to the pelvic bone, with 2 µl/injection at the circumferential left, circumferential right, and midline locations. The bladder wall injections (2 µl/ injection) were performed on each side of the bladder dome. All injections were delivered with a 30-gauge needle (HAMC7803-07) attached to a Hamilton 5 ml syringe (HAMC7634-01). Mice were housed individually and allowed to recover for 1 week before undergoing transcardial perfusion. Following complete perfusion, the spinal cord and brain were collected. The spinal cord below the L2 vertebra was isolated, as it contains the levels L5-S1 relevant to colon and bladder sensory processing (11).

### Fluorescence In Situ Hybridization and Immunohistochemistry

Cryosections (20 µm) from the spinal cord and brain, fixed in 4% paraformaldehyde, were used for both fluorescent ISH and immunostaining. Fluorescent ISH was performed using RNAScope (ACD, Multiplex Fluorescent V2) according to the manufacturer’s protocol using the following probes: *Slc32a1*/*Vgat* (319191), *Slc17a6/Vglut2* (319171-C3), *Lhx1* (488581), *Crh* (316091-C4), *Fos* (316921-C2). Immunohistochemistry was performed using the following primary antibodies: Tyrosine hydroxylase (Millipore-Sigma, MAB318, and ImmunoStar, 22941), cFOS (Santa Cruz, sc-52, and Cell Signaling, 2250), RFP (Rockland Immunochemicals, 600-401-379S), WGA (Millipore-Sigma, T4144), GFP (AvesLabs, GFP-1010). For detection, sections were labeled with species-appropriate Alexa Fluor-conjugated secondary antibodies (Invitrogen, #A-21447, #A10042, #A-11055, #A-21121, #A-11001; 1:1000 dilution). A complete list of antibodies and dilutions used in immunohistochemical analyses is provided in Table S2.

Slides were imaged with a Keyence BZ-X800 widefield fluorescence microscope or autoscanned using an epifluorescence ZEISS Axioscan 7 Slide Scanner (Carl Zeiss Microscopy, Germany). Confocal images were acquired at the UCSF Center for Advanced Light Microscopy (CALM) using a CSU-W1 Spinning Disk/ High Speed Widefield with an Andor Zyla sCMOS camera and MicroManager v2.0gamma, or a confocal laser scanning microscope (Leica TCS SP8X, Germany). Images were processed and quantified using ImageJ Fiji (NIH) and the Cell Counter plugin v2, LAS-Lite (Leica), ZEN BLUE (Carl Zeiss Microscopy), and Corel-DRAW Graphic Suite software. Three representative views of each sample were selected.

### Quantification of Colon and Bladder Co-Activation

The number of tdTomato-positive, cFOS-positive, and co-labeled (tdTomato/cFOS) neurons was counted in regions of interest (single-plane images) using the cell detection tool in QuPath (version 7.0). Neuronal counts were obtained from 5-10 sections/spinal cord/mouse and 1-3 sections/BAR/LC or PAG per mouse. Only cells with a neuronal morphological profile and intact nuclei (identified by NucBlue) were included in the counts.

### Statistics

Statistical tests were performed using Prism 11.0.2 (GraphPad). Multiple comparisons correction for One-way, Two-way, and repeated-measures (RM) ANOVA was performed using the post hoc tests indicated in each figure legend. For all panels in the Main and Extended Data Figures, N = biological sample size, and n = replicate count. Outliers were identified by Grubbs test (Alpha = 0.05). Unless otherwise noted, data are presented as mean ± SEM or as paired or unpaired scattered plots. Sample sizes are based on prior work from our labs; however, no specific statistical calculation was performed to determine them. For AAV viral vector injections, mice of identical genotypes were randomly selected from littermate pools to receive functional or control viral vectors. Experimenters were blinded to the type of AAV received/genotype of the mice under study during experimental analyses. For cFOS quantification in bladder/colon distensions, unpaired t-tests and One-Way ANOVA with Bonferroni multiple comparison tests were used for parametric distributed data, One-way Kruskal-Wallis test with Dunn’s multiple comparisons tests were used for non-parametric data. A description of the test and results is provided in Table S3.

## ACKNOWLEDGMENTS

We thank all members of the Ingraham group for helpful suggestions, critical comments, and strategic contributions, including David Julius, Kevin Yackle, Wendy Yue, and Sean Li for their pharmacological insights and intellectual engagement in this project. We especially acknowledge early intellectual and technical input from William Krause, as well as assistance from Yicen J. Zheng. We also thank Drs. TA Wang and LY Jan for sharing the optimized WTR toolkit prior to publication. This work was supported by grants from the US NIH, including K99DK139516-01 (AV), R21AG086774, R01DK135714-01, R01DK147593-01 (HAI), U01NS136405, R01EY030138 (XD), R01 ES035020 (KPKS), funding from the Australian Research Council, Discovery Project DP220101269 (AMH and SMB) and the National Health and Medical Research Council of Australia (NHMRC) with Investigator Leadership Grant APP2008727 (SMB). Core services were made possible by funding from the NIH NIDDK Support for the Wisconsin O’Brien Center for Benign Urologic Research, Rodent Urinary Function Testing Core, U54 DK104310 (KPKS). We acknowledge the Center for Advanced Light Microscopy (CALM) at UCSF, where data were acquired on a CSU-W1 Confocal obtained using NIH S10 Shared Instrumentation grant (1S10OD017993-01A1).

## DATA AVAILABILITY STATEMENT

All data generated or analyzed during this study will be included in the published article (and its supplementary information files). Supplementary Figures and Movies will be distributed upon request.

## CONTRIBUTIONS

LECZ, AV, KKS, SMB, AMH, and HAI conceived and designed the experiments, interpreted the results, and wrote the paper. LECZ designed and conducted all stereotaxic surgeries and viral vector injections, which were followed up by histological confirmation. He also conducted all VSA and activity assays. KKS performed all in vivo cystometry and uroflow assays. AMH conducted retrograde tracing into the spinal cord and cFOS analyses following colon/ bladder distensions. AV performed surgical implantation of wireless transmitters for visceral motor reflex (VMR) assays and colonic transit. FCN aided in blood collection. KG assisted with the quantification of histological results. XD and JH provided mWmC and WTR viral vectors, as well as guidance on the studies.

Note that we requested an external review of this study by **QED Science** (https://www.qedscience.com) after uploading to bioRxiv, a program that provides a score based on novelty as well as experimental gaps. This study outperformed 99% of >50,000 papers based on their calculated benchmarks.

